# Hijacking of Cellular Functions by Severe Acute Respiratory Syndrome Coronavirus-2. Permeabilization and Polarization of the Host Lipid Membrane by Viroporins

**DOI:** 10.1101/2022.04.14.488372

**Authors:** Emmanuelle Bignon, Marco Marazzi, Antonio Monari

**Affiliations:** Université de Lorraine and CNRS, UMR 7019 LPCT, F-54000 Nancy, France; Universidad de Alcalá, Departamento de Química Analítica, Química Física e Ingeniería Química, Grupo de Reactividad y Estructura Molecular (RESMOL), Alcalá de Henares, Madrid, Spain; Universidad de Alcalá, Instituto de Investigación Química “Andrés M. del Río” (IQAR), Alcalá de Henares, Madrid, Spain; Université Paris Cité and CNRS, ITODYS, F-75006 Paris, France

**Keywords:** Molecular Dynamics, Coronaviruses, Free Energy Methods, Ion Channels, Cellular Membrane, Viral Inflammasome

## Abstract

As all viral infections, SARS-CoV-2 acts at multiple levels hijacking fundamental cellular functions and assuring its replication and immune system evasion. In particular, it has been observed that the viral 3’ Open Reading Frame (ORF3a) codes for a hydrophobic protein which embeds in the cellular membrane, where it acts as an ion viroporin and is related to strong inflammatory response. Here we report equilibrium and enhanced sampling molecular dynamic simulation of the SARS-CoV-2 ORF3a in a model lipid bilayer, showing how the protein permeabilizes the lipid membrane, via the formation of a water channel, which in turn assures ion transport. We report the free energy profile for both K^+^ and Cl^-^ transfer from the cytosol to the extracellular domain. The important role of ORF3a in the viral cycle, and its highly conservation among coronaviruses, may also make it a target of choice for future antiviral development, further justifying the elucidation of its mechanism at the atomistic level.

**TOC GRAPHICS:** 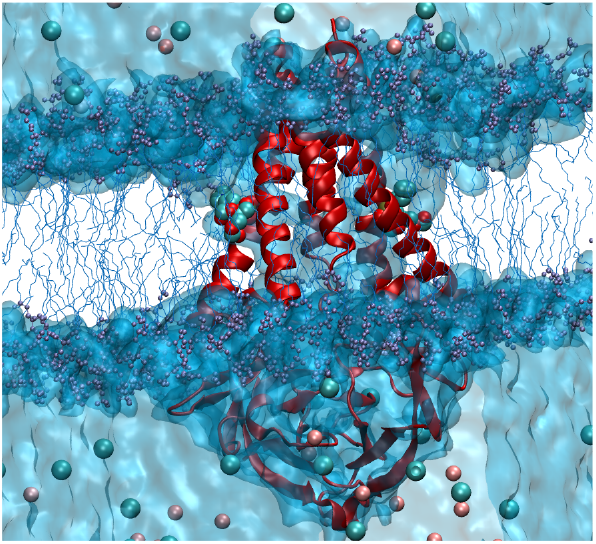

---

A novel Betacoronavirus, firstly identified in Wuhan China^1^ at the end of 2019 and named SARS-CoV-2,^2^ is the causative agent of the COVID-19 pandemics,^3^ which at the beginning of 2022 is still ravaging worldwide. Because of SARS-CoV-2 high contagiousness, and the possible emergence of severe infection, especially in elder subjects or in patients presenting comorbidity, COVID-19 is causing an important threat on Health and Social systems, notably requiring the implementation of severe social distancing measures.^4^ While the development of effective vaccines, also based on the novel messenger RNA (mRNA) approach,^5–9^ has significatively contributed to limit COVID-19 impact, the development of novel antivirals is still crucial, especially in case of immunodeficient patients or to counteract possible vaccine-resistant variants.^10^ In this respect, targeting highly conserved proteins in SARS-CoV-2, and more generally in coronavirus strains, is highly suitable.

As other parent coronaviruses, SARS-CoV-2 is an enveloped positive-sense RNA virus,^11^ encoding a rather large genome of about 30,000 nucleobases. The analysis of SARS-CoV-2 genome reveals the presence, in addition to a 5’ untranslated terminal region (UTR),^12^ of different areas leading to the expression of large non-structural proteins,^13^ mainly exerting enzymatic functions necessary to the viral reproduction and maintenance. More in detail, the native viral polyprotein is self-cleaved by the two proteases (3CL-like and papain-like)^14–16^ to finalize the maturation of all its non-structural proteins. Furthermore, other structural proteins necessary to ensure the structuration of the virion are also expressed, including the well-known Spike protein,^17,18^ which protrudes from the viral capsid and allows the entry of the virus into the host cells. The following region of the viral genome, the 3’ Open Reading Frame (ORF3a) is, instead, associated with the encoding of hydrophobic proteins which may get involved in vesicle formation.^19,20^

As a matter of fact, Spike protein,^21^ together with proteases^22^ and to a lesser extent RNA-dependent RNA polymerase (RdRp),^23^ represent the target of choice for both vaccine^24^ and antiviral development. However, it has recently been pointed out that ORF3a encodes a membrane protein which acts a viroporin,^25–28^ inducing cellular membrane permeation and depolarization, probably acting as a cation (mostly K^+^) channel. Indeed, viroporins,^29,30^ although not strictly necessary for viral maturation, may favor the virus replication and diffusion contributing to its infectivity.

Furthermore, their potential to permeabilize membranes have also been associated with the development of a strong cytokine-based inflammatory response,^25^ which is usually the cause of serious outcomes in COVID-19 patients, especially in presence of inflammatory related comorbidity such as diabetes or obesity.^31^ Finally, ORF3a appears highly conserved between the different coronaviruses and the SARS-CoV-2 mutants.^28^ Hence, blocking its function by specific ligands could represent a most valuable drug-design strategy.

The structure of ORF3a embedded in lipid nanodiscs has been recently resolved by cryo-electron microscopy (Cryo-EM),^27^ revealing that the functional unit is represented by a homodimer (Figure 1A), although a loosely bound tetramer is also found in infected cells. The transmembrane domain of each monomer is constituted by three α-helices which intertwine to provide a central channel flanked by polar residues. The cytosolic region, on the other hand, forms a bulge and is highly structured by β-sheets domains. In vitro measurements^27^ have also shown that ORF3a may act as an ion channel with a slightly preferred selectivity for cations, especially K^+^, over anions, while it is inhibited by polyamines. If the porin activity of ORF3a may be supported by the presence of the central polar channel at the monomers interface, the actual mechanisms of membrane permeabilization and polarization is still debated.^27^ In particular, it is still unclear whether water conductivity should take place entirely through the central channel, albeit requiring consistent structural reorganization, i.e. a situation reminiscent of cellular voltage-^32^ or ligand-gated^33^ ion channels, or laterally through polar grooves developing along the α-helices.

**Figure 1.**
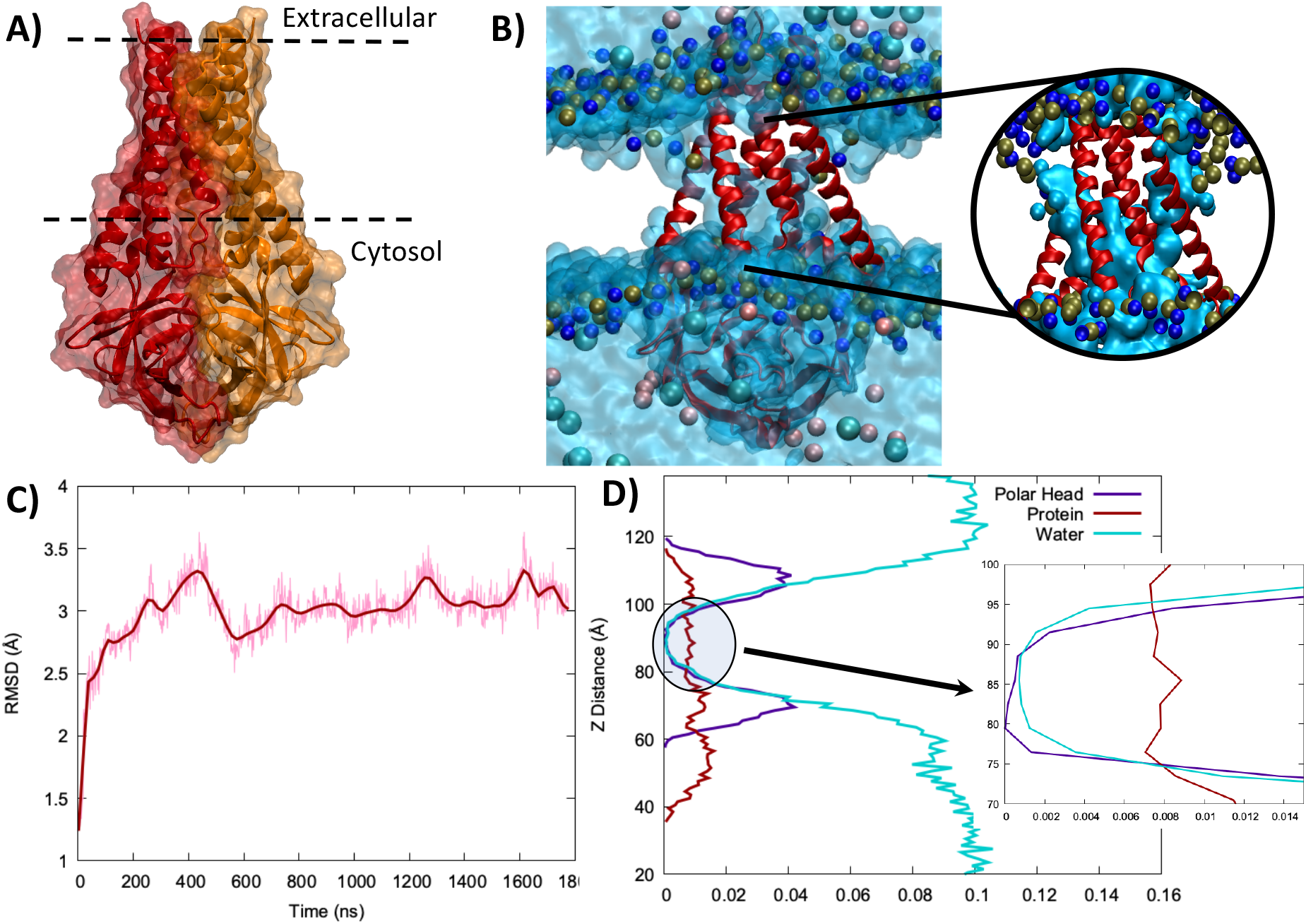
A) Structure of the ORF3a dimer: the two monomers are represented in red and orange, respectively. The position of the membrane polar heads is also schematically indicated as well as the cytosolic and extracellular sites. B) Representative snapshot extracted from the MD simulation of ORF3a embedded in a lipid bilayer. Note that K^+^ and Cl^-^ ions are represented as van der Waals spheres, in pink and cyan, respectively, while the polar heads are depicted as blue and bronze spheres. The water is depicted as surfaces. In the inlay a zoom of the transmembrane region of ORF3a is provided with water in contact to the protein represented as a solid surface to highlight the lateral polarization channel. C) Time series of ORF3a RMSD and D) density profile along the membrane axis of ORF3a, water, and polar heads.

In this work, we report classical molecular dynamics (MD) simulations exceeding the μs timescale and coupled with free energy methods to unravel the structural and dynamical features of ORF3a embedded in a model lipid membrane, and to assess the thermodynamic profile for K^+^ and Cl^-^ transfer from the cytosolic to the extracellular domain. The ORF3a initial structure was obtained from the one resolved by Kern et al.^27^ (pdb code 6XDC), embedded in a double layer of 1-palmitoyl-2-oleoyl-sn-glycero-3-phosphocholine (POPC) lipids, and further solvated with water buffer including physiological salt (KCl) concentration of 0.15 M.^34–36^ The initial system was prepared using CHARMM-GUI input generator, after modeling the 175-180 missing loop with the SwissModel server. Both equilibrium and free-energy MD simulations have been performed using NAMD^37,38^ and analyzed with VMD.^39^ Amber14 force field for both protein and lipid was consistently used, isobaric and isothermal conditions (NPT) have been applied, and the Newton’s equations of motion have been integrated with a time step of 4 fs using Hydrogen Mass Repartition (HMR)^40^ in combination with Rattle and Shake.^41^ Protein-lipid interactions were calculated by analyzing the trajectory data through the ProLint software, freely available by the *prolintpy* package.^42^ All protein residues and a cutoff of 10.0 Å distance from each residue to any membrane lipid was considered. Extended Adaptative Biased Force (eABF),^43,44^ as implemented in Colvar^45^ and NAMD,^37,38^ has been used to calculate the free energy profiles for K^+^ and Cl^-^ penetration. More details on the computational methodology and strategy are provided in Electronic Supplementary Information (ESI).

Following the equilibration of both the protein and the lipid bilayer, ORF3a experiences a remarkable stability as shown in Figure 1C. Indeed, the Root Mean Square Deviation (RMSD) of the protein plateaus at around 2.96 ± 0.28 Å. All the main features identified in the Cryo-EM experimental structure are maintained in our simulations. In particular, the organization of the transmembrane domain, and the α-helical regions appear as particularly rigid, as compared to the cytosolic domain, which experiences a higher flexibility. More importantly, the visual analysis of the MD simulation already confirms that ORF3a is indeed permeabilizing the lipid bilayer inducing a transmembrane water channel (Figure 1B), which in turn may also lead to ion conductivity. The permeabilization is also confirmed by the density profile along the membrane axis (Figure 1D), which shows that water density does not go to zero even at the hydrophobic center of the membrane. The permeabilization of the membrane by ORF3a proceeds via the hydration of two lateral polar channels (constituted mainly by tyrosine and leucine residues) flanking the α-helices. These two channels converge towards two gates buried in the membrane hydrophobic core and constituted by positively charged arginine and lysine residues, which, in turn, drive the water channels inside the central pocket and further into the cytosolic domain. The cytosolic side, as well as the central pocket, are largely composed of polar or charged residues, hence facilitating water exchange (see Figure S1).

The charged or polar amino acids on the a-helix surface may even exhibit transient, yet relatively-long lived, interactions with the polar heads on the extracellular side as shown in Figure 2, where the time series of the distance between arginine and lysine residues with the closest POPC phosphate groups are reported. This occurrence leads to a further curvature of the lipid membrane around ORF3a, thus facilitating the emergence of the later water channels. However, this interaction is not essential for the membrane permeabilization (Figure 2 A) as also supported by the asymmetry of the interaction, which is stronger for N357, N320, and K61, i.e. on one of the channels, than for N82, N119, and K299. The latter interaction is, indeed, completely lost at around 1 μs, resulting in the release of the lipid polar head without compromising water conductivity (Figure 2 C). Furthermore, breaking the residual interaction of the closest polar head lipid with N357, N320, and K61 through Steered MD (STMD, see details in ESI) to position the phosphatidyl choline moiety back in the polar head region results in neither the destabilization of the global protein-water interface nor in the loss of permeabilization – see Figure S2.

**Figure 2.**
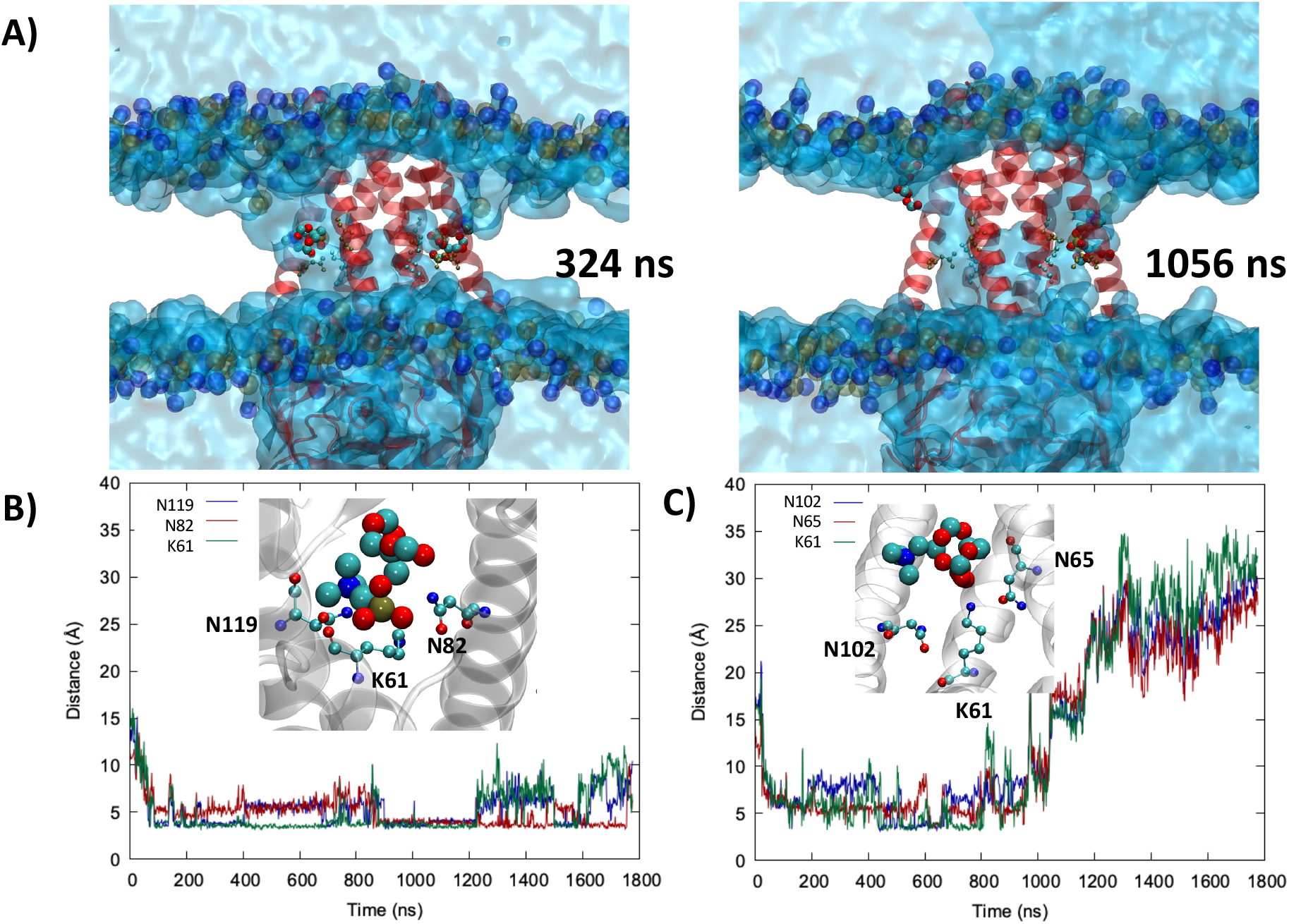
A) Two representative snapshots of the ORF3a dynamics showing the interaction between the lipids with arginine and lysine residues at early (324 ns) and late (1056 ns) stages of the MD simulation showing the release of one of the lipids while the water channels remain unaffected. The displaced POPC unit is represented by van der Waals representation, while the interacting amino acid by CPK drawing. B) Time series of the distances between the lateral chain nitrogens of K61, N82 (chain B), and N119 (chain B) and the phosphate atom of the closest POPC and C) between the lateral chain nitrogens of K61 (chain B), N65, and N102 and the phosphate atom of the closest POPC. The specific interactions between the polar and charged amino acid and the polar head are depicted in the inlay.

When focusing on protein-membrane interactions, we evidence a clear asymmetry between extracellular-side and cytosol-side gates. More specifically, a consistent number of residues (highlighted in yellow in Figure 3 A) experiences a relatively long contact time with at least one polar head of the membrane surface. Such contact time is given in terms of percentage referred to the whole production trajectory (Figure 3 B). On the other hand, only two residues form a stable interaction with extracellular-side lipid polar heads (highlighted in red in Figure 3 A), hence supporting a much higher lipids mobility at the extracellular gate. This observation is also partly justified by the overall smaller surface of the extracellular gate, especially when compared to the larger and well stabilized cytosol gate, which is constituted by almost all the residues belonging to the terminal sides of the six transmembrane α-helices and pointing toward the cytosol. The two protein monomers (highlighted in purple and green in Figure 3 C) both contribute to these type of interactions, although not equivalently, with the most relevant residues, i.e. those showing a contact mean duration greater than 13%, being polar (S135), charged (R68), amphipathic (Y141, P159) but also hydrophobic (L129, W69 (chain B), L140 (chain B), A144 (chain B)). Although the chosen cut off of 13% contact is largely arbitrary it allows to identify the different specific interactions taking place.

**Figure 3.**
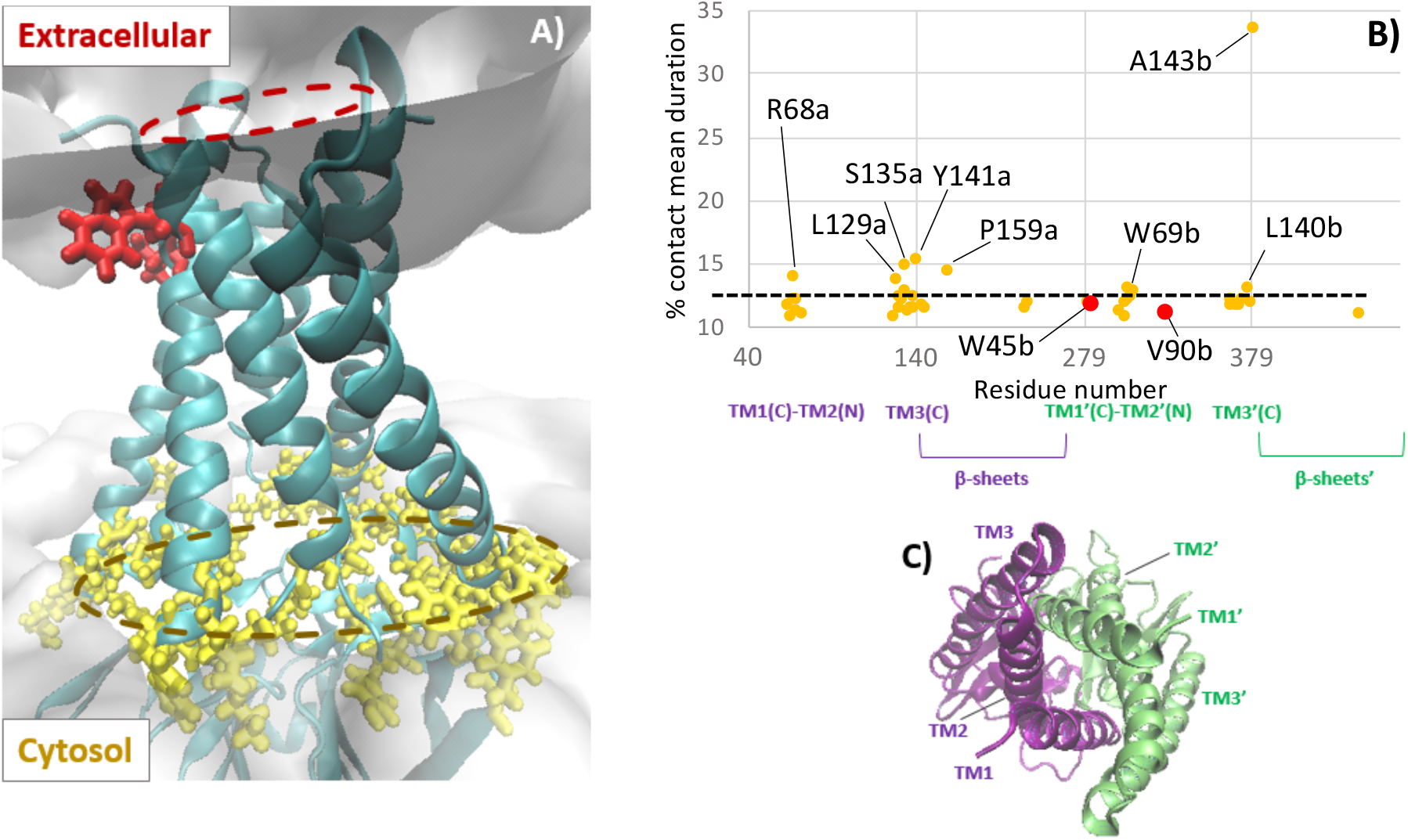
A) ORF3a dimer representation, within which the residues forming stable protein-lipid interactions on the cytosol-side (yellow) and on the extracellular-side (red) have been highlighted. B) Protein-lipid contacts shown as the contact mean duration experienced by each residue along the trajectory, following the same color code. All residues with contact mean duration greater than 13% are highlighted, as well as the two residues on the extracellular side, the labels a and b are postponed to the aminoacid one-letter code to differentiated between the two monomers. C) Top view of ORF3a (extracellular side), with each monomer depicted in purple and green.

Interestingly, in the course of the equilibrium MD simulation we have observed multiple occurrences of a transient binding of Cl^-^ ions inside the ORF3a pocket (Figure 4). The ions experienced variable persistency time inside the pocket, up to 500 ns (Figure 4 B). For all the observed cases the penetration took place from the cytosolic region through a gate involving the polar N144, N161, T64, and K66 (Figure 4 C). Quite interestingly, we have also evidenced that the ORF3a pocket can indeed accommodate, despite its relatively small volume, two anions.

**Figure 4.**
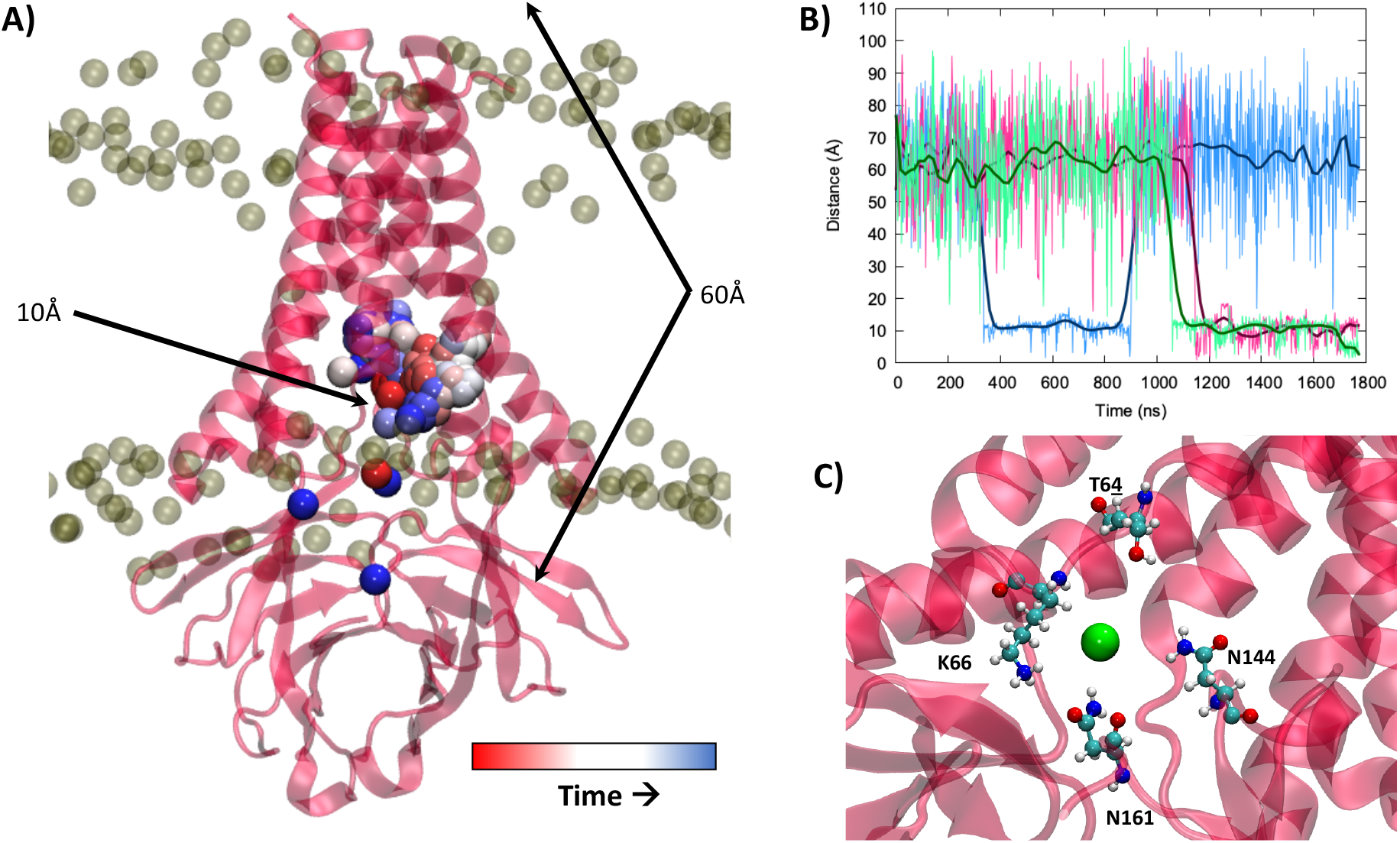
A) Evolution of the position of the Cl^-^ ion entering the pocket in the non-biased MD simulation. The ion is represented as a van der Waals sphere whose color code indicates the time. B) Time series of the distance between three Cl^-^ anions entering the ORF3a pocket and the center of the protein, the positions corresponding to 10 and 60 Å distance from the center of the membrane are also indicated on panel A. C) Entry gate for the Cl^-^ ion (green van der Waals sphere), the surrounding polar amino acids are represented in CPK.

While we have observed the spontaneous anion release back to the cytosol, no occurrence of ion transfer to the extracellular medium was observed in the course of the equilibrium MD simulations. Thus, we resorted to enhanced sampling methods (eABF,^46^ see details in ESI) to identify the free energy profiles related to both K^+^ and Cl^-^ transfer using the distance from the center of mass of the membrane (taken as Q57, S58 from both monomers, a lock invoked in the literature)^27^ projected on the Z membrane axis as the collective variable.

Both K^+^ and Cl^-^ follow the same path for their transfer through the membrane (Figure 5 A). First, the ions move up within the central cavity of ORF3a from the cytosolic side. Then, they pass over the Q57-S58 lock and slide towards the lateral channel. This path is consistent with the water channel described for unbiased simulations. The K^+^ trajectory exhibits a slight increase of the free energy upon the entrance of the ion in the cavity on the cytosolic side, whereas Cl^-^ reaches a minimum energy state when arriving at the top of this cavity (Z distance ~4 Å, see Figure 5 B). Indeed, Cl^-^ makes stabilizing interactions with S60, H78 and the Q57-S58 lock of both ORF3a monomers, prior to overpassing the latter. However, both K^+^ and Cl^-^ have to cross a ~7-8 kcal/mol energy barrier to achieve the transfer from the cytosol to the extracellular side. Interestingly, while the K^+^ transfer energy peaks at Z distance = 0 Å (passing the Q57-S58 lock), the Cl^-^ transition state occurs slightly later along the collective variable, at a distance of 7 Å, corresponding to the hydrophobic A54, T89 (chain B), Q116 (chain B) residues – Figures 5 C-E and S3. In both cases, a drop in the free energy of ~3 kcal/mol is observed after bypassing the energy barrier. Overall, albeit the slight shift due to their opposite charge leading to contrasted interaction patterns, both K^+^ and Cl^-^ show an easy transition from the cytosol to the extra-cellular medium. The differences observed, especially in the initial portion of the collective variable between the cation and the anion are also coherent with the equilibrium MD simulation, which has provided multiple occurrences of Cl^-^ penetration inside the ORF3a cavity. Furthermore, the presence of an initial free energy plateau for K^+^ penetration may participate in lowering the activation energy for the rate-determining step, hence justifying the slight selectivity for this ion. In contrast, the free energy minimum observed inside the cavity for chlorine may lead to a trap significantly increasing its residence time. Interestingly, the amino acids identified in the channel previously predicted by Kern et al. match with the path of the ions in our eABF simulations (see Figure S4).

**Figure 5.**
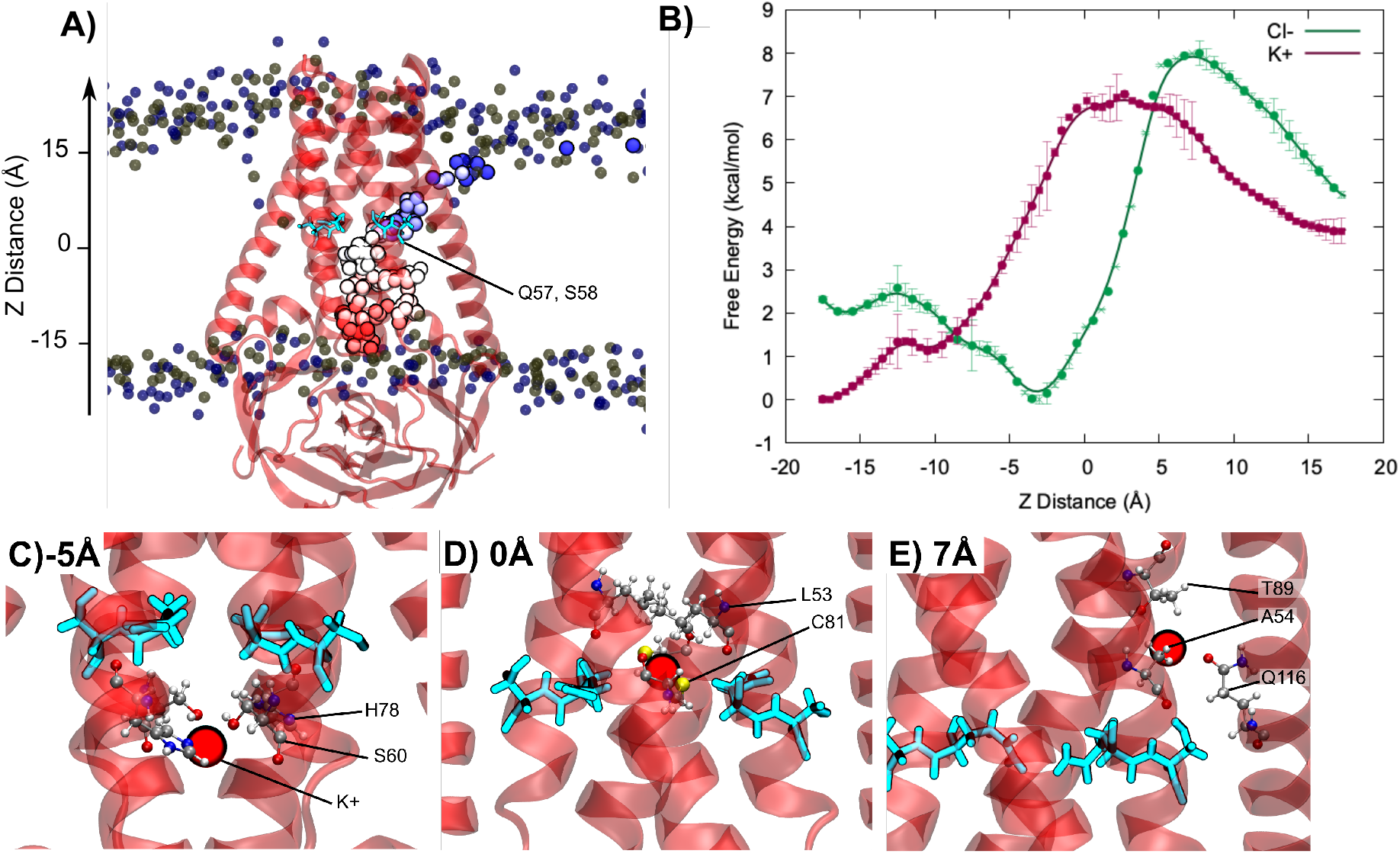
A) Potassium ion transfer path through ORF3a. The Z distance collective variable is defined as the projection of the distance between the ion and the center of mass of the Q57 and S58 gate residues (depicted in cyan) from each monomer on the Z axis. The ion is colored according to the trajectory time (from red to blue). B) Free energy profiles of K^+^ (red) and Cl- (green) transfer from the cytosol to the extracellular medium, the statistical uncertainties are also reported as error bar as well as in Figure S4. C) Interactions of K^+^ with each monomer’s S60 and H78 at the entry gate (−5 Å, cytosolic side), D) with Q57, S58, C81 and L53 at the center of the membrane (0 Å), and E) with A54, T89 (chain B), Q116 (chain B) at the exit gate (8 Å, extracellular side).

Our simulations have provided a clear vision of the ORF3a permeabilization of a model host lipid membrane. More specifically we have clearly evidenced the spontaneous formation of a water channel. the most favorable permeabilization pattern involves the presence of lateral hydrophilic channels, which are constantly active and allow the transfer of both solvent and ions that favors the transport of ions. This mechanism has been clearly identified as the most favorable one, especially compared to the activation of a central channel which should require a large conformational reorganization. Such less-common ion permeabilization was indeed found to be the most feasible both kinetically, since it does not require channel activation/deactivation cycles, and thermodynamically, since it surely requires the smallest possible energy barrier to be overcome, as previously suggested in some other systems.^47,48^ In particular, the free energy profiles for the ions transfer through the transmembrane protein, evidence its feasibility for both positively and negatively charged species, while highlighting the importance of the Q57-S58 lock for the K^+^transition from the cytosol to the extracellular domain and revealing a slightly different profile for Cl^-^.

Apart from ions, we would also like to draw attention on the presence, in the beginning of the simulation, of a lipid molecule between two α-helices (sketched in yellow and red in Figure S5), still partially attached to its original extracellular membrane side. Interestingly, by analyzing the distance of such outbound lipid from the edges of a transmembrane α-helix (R68-A99), it can be seen how the lipid is transferred across the membrane without any external bias, to almost reach the cytosol membrane side finally. This indication mildly suggests that the eventual ORF3a central channel might be accessible for lipid translocation, due to the presence of a hydrophobic cavity shielding the lipid molecule from the aqueous environment (see also Figure 2A). Nevertheless, more realistic membrane compositions should be used to confirm the eventual co-role of ORF3a as lipid transporter.

All in all, our results allow to characterize, at an atomistic level, the mechanisms underlying permeabilization and polarization of the membrane by SARS-CoV-2 ORF3a viroporin-like protein. In this sense, and due to the important role of ORF3a for the viral life cycle, they could offer original drug design strategy aimed at blocking the transfer gates by the interaction with specific drugs. The high conservation of ORF3a among coronaviruses further enhances the importance of this study and the soundness of the envisaged drug-design strategy, while providing the elucidation of a key, and often overlooked, mechanism of SARS-CoV-2 cycle.

## ASSOCIATED CONTENT

### Supporting Information

The following files are available free of charge.

Supplementary text and figures (PDF file)

Movie of the Cl-transfer through ORF3a (MPG file)

## AUTHOR INFORMATION

### Notes

The authors declare no competing financial interests.

## ACKNOWLEDGMENT

The authors thank GENCI and Explor computing centers for computational resources. E.B. thanks the CNRS and French Ministry of Higher Education Research and Innovation (MESRI) for her postdoc fellowship under the GAVO program. A.M. thanks ANR and CGI for their financial support of this work through Labex SEAM ANR 11 LABX 086, ANR 11 IDEX 05 02. The support of the IdEx “Université Paris 2019” ANR-18-IDEX-0001 and of the Platform P3MB is gratefully acknowledged. M.M. and A.M. acknowledge funding from DISCOVER-UAH-CM project, cofounded by Community of Madrid (CAM) and European Union (EU), through the European Regional Development Fund (ERDF) and supported as part of the EU’s response to COVID-19 pandemic.

