## Supplementary Information for "Hijacking of Cellular Functions by Severe Acute Respiratory Syndrome Coronavirus-2. Permeabilization and Polarization of the Host Lipid Membrane by Viroporins"

### SUPPLEMENTARY COMPUTATIONAL METHODS

*Steered Molecular Dynamics*

A steered Molecular Dynamics run was performed in order to evaluate the effect of the buried lipid polar head on the permeabilization of the membrane by ORF3a. The phosphatidyl choline moiety was pull towards the membrane polar heads region on the extracellular side, by imposing a strong constraint (150 kcal/mol) on the projection of the distance between the lipid (atoms 7429-7466) and the center of mass of the first ORF3a monomer (atoms 1-3222) on the Z axis. This Z distance was increased from 17 Å to 31 Å in 800 ps. The final structure was subsequently relaxed along a 550 ns unbiased MD run – see Figure S2.

*Free energy calculations*

The potential of mean force (PMF) of the transfer of K+ or Cl- ions from the cytosol to the extracellular medium was obtained using the extended adaptative biasing force (eABF) technique. In each case, the ion (residue 1278) was pull through ORF3a along the projection of its distance to the center of mass of the Q57-S58 lock on the Z axis, from -17.5 Å to +17.5 Å. The spring stiffness and the oscillation period of the fictitious particle were set to 0.5 Å and 100 fs, respectively. The path was divided into 8 windows of 5 Å, and upper and lower walls were set at -0.5 and +0.5 of the boundaries with a 50 kcal/mol force constant. Gaussians with a width of 0.1 were deposited with a height of 5 Å and a weight of 0.1 kcal/mol. Each window was sampled for 180 ns and output values were dumped every 1000 steps (4 ps).

**SUPPLEMENTARY FIGURES**

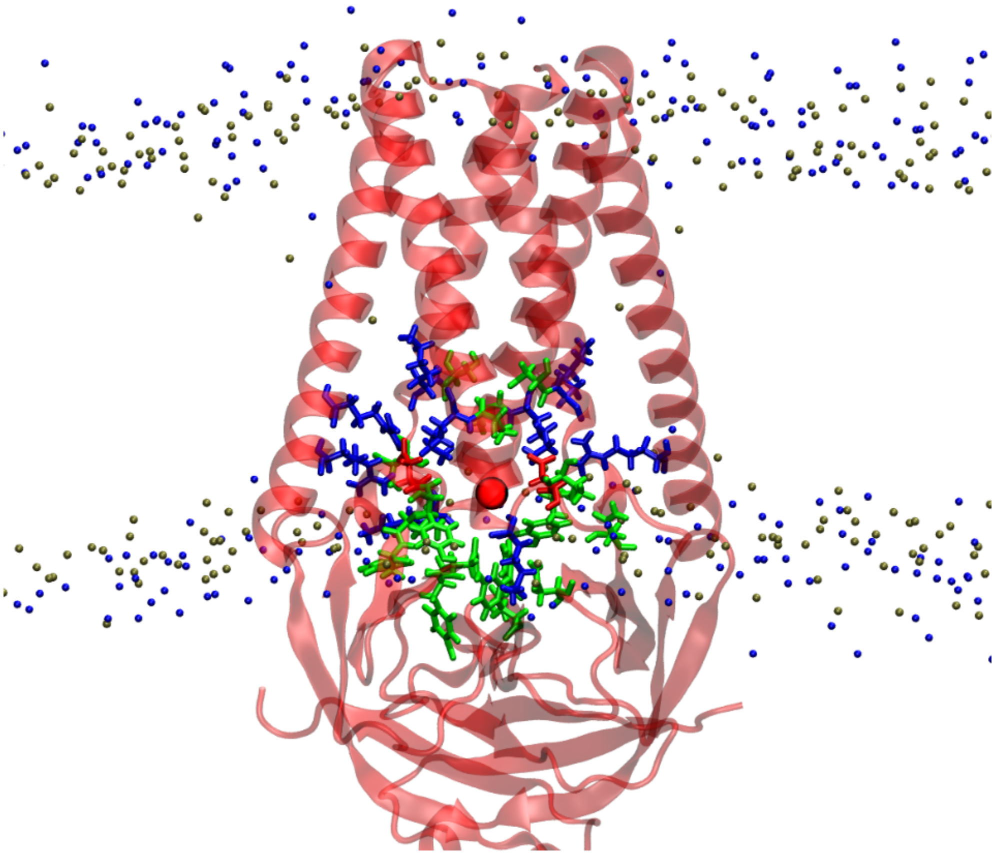

Figure S1. Location of polar, basic and acidic amino acids on the cytosolic side and within the central pocket, represented in green, blue and red, respectively.

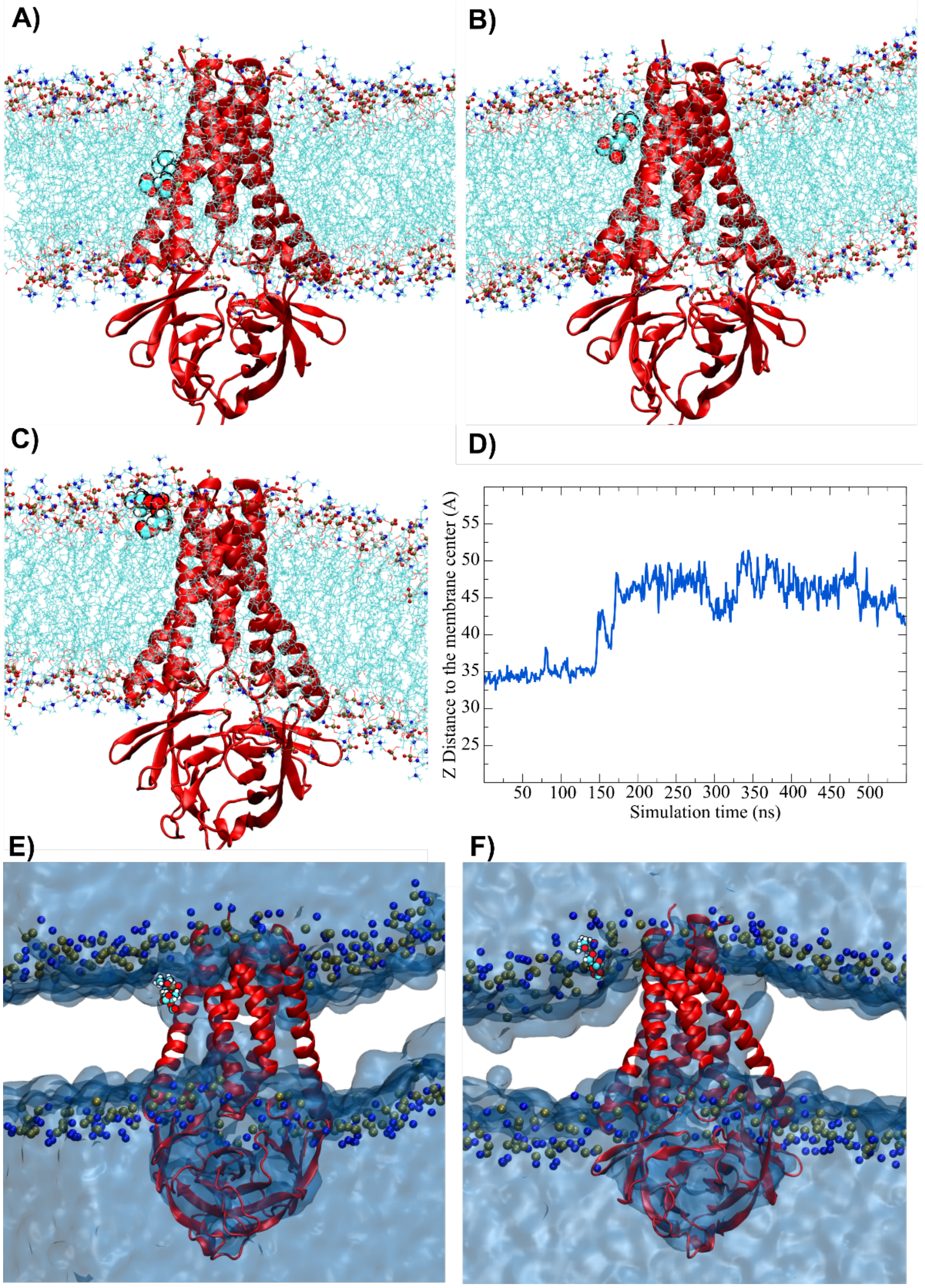


Figure S2. Position of the displaced lipid A) before MD, and B) after the steered MD run. C) Structure of the complex and location of the displaced after 550 ns of relaxation following the STMD and D) evolution of the Z distance as used as collective variable for the STMD along the unbiased final run. E) Conserved water flooding of the channel after STMD and F) after the unbiased run.

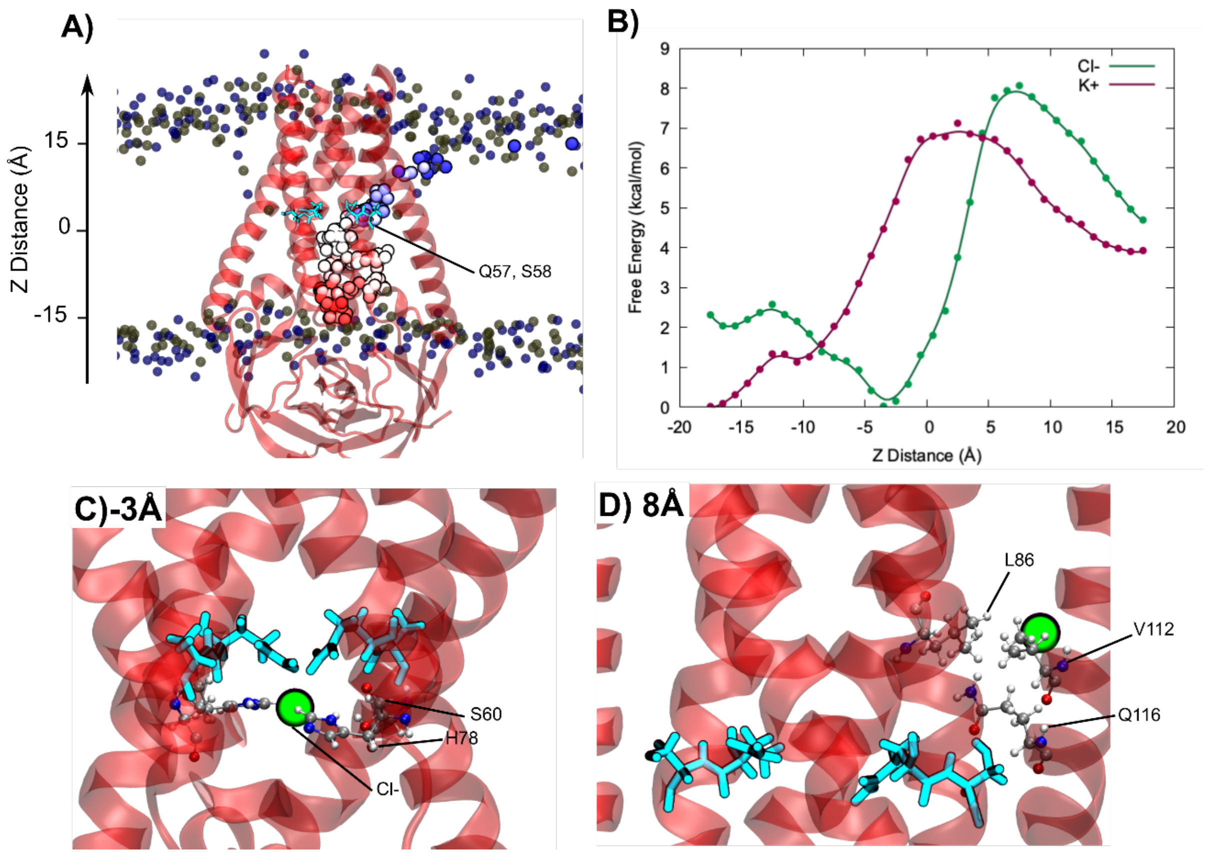


Figure S3. A) Chloride ion transfer path through ORF3a. The Z distance collective variable is defined as the projection of the distance between the ion and the center of mass of the Q57 and S58 gate residues (depicted in cyan) from each monomer on the Z axis. The ion is colored according to the trajectory time (from red to blue). B) Free energy profiles of K+ (red) and Cl- (green) transfer from the cytosol to the extracellular medium. C) Interactions of Cl- with each monomer's S60 and H78 below the Q57-S58 lock (-3 Å, cytosolic side, minimum in the energy path), and D) with L86, V112, and Q116 right after passing the lateral exit gate (8 Å, extracellular side).

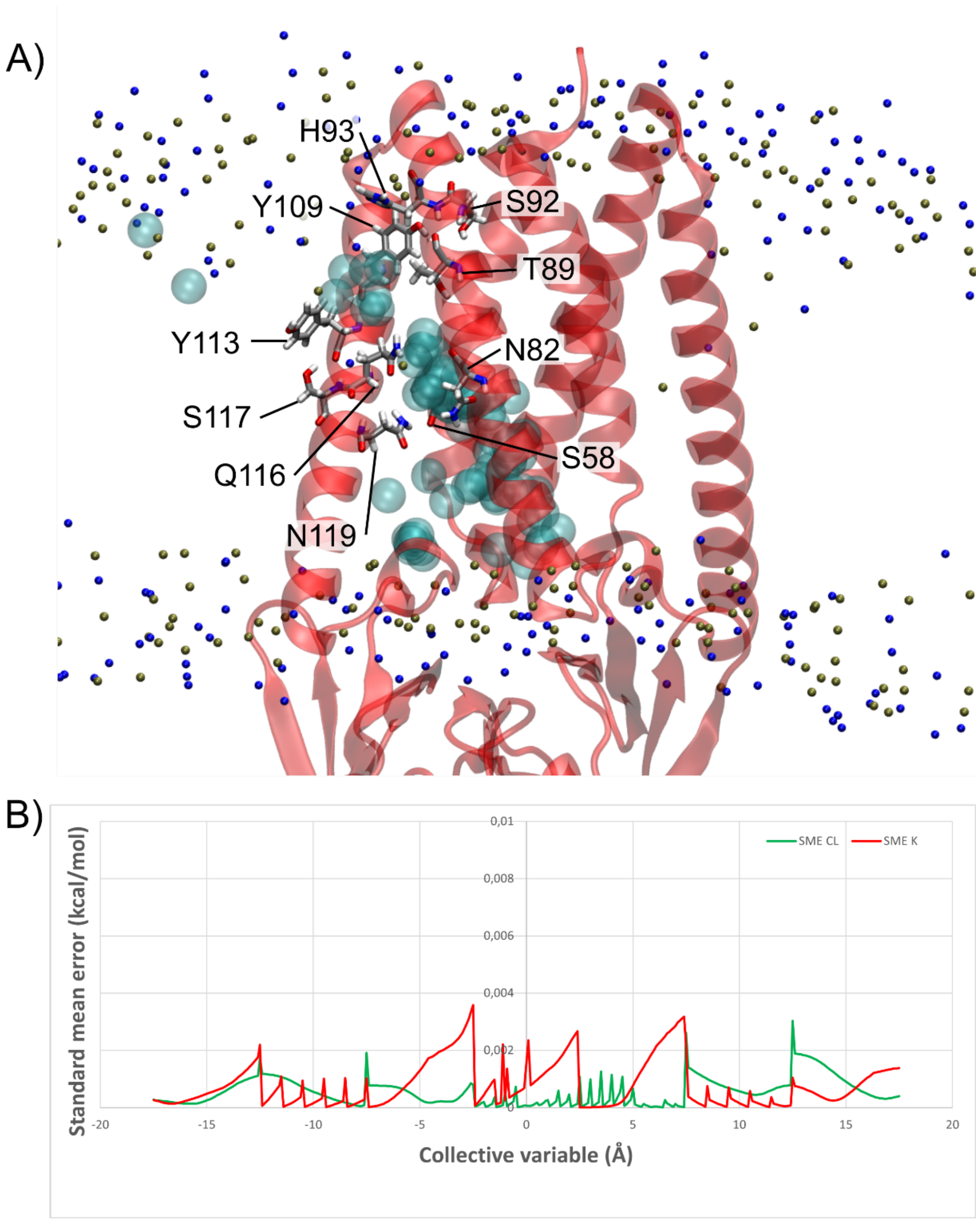


Figure S4. A) Amino acids along the ion path through ORF3a as hypothesized by Kern et al., highlighting the very good agreement with the results our from free energy calculations. The path is depicted in transparent blue, and amino acid in licorice. B) Histograms of standard mean errors calculated from eABF calculation for both the chloride (green) and the potassium (red) ions.

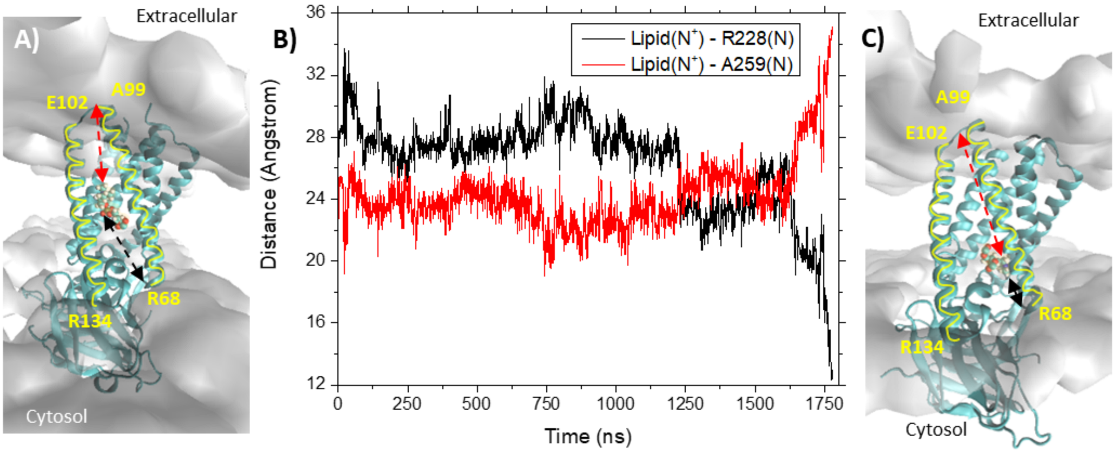


Figure S5. Initial A) and final C) position of a lipid within ORF3a (polar head shown in CPK and highlighted in light orange). Two α-helices, E102-R134 and R68-A99, along which the lipid transport is observed, are sketched in yellow. The membrane polar heads constituting the extracellular and cytosol sides are also shown in transparent grey. The distance between the N+ of the lipid and the N of A99 or the N of R68 (red and black dotted arrow, respectively) is also shown. B) The same color code (red and black) is used to show such distances along the trajectory, highlighting the transfer of the lipid in the final part of the trajectory (from ca. 1600 ns).
